# Effects of water nutrient concentrations on stream macroinvertebrate community stoichiometry: a large-scale study

**DOI:** 10.1101/2024.02.01.574823

**Authors:** M. Beck, E. Billoir, P. Usseglio-Polatera, A. Meyer, E. Gautreau, M. Danger

## Abstract

Basal resources generally mirror environmental nutrient concentrations in the elemental composition of their tissue, meaning that nutrient alterations can directly reach consumer level. An increased nutrient content (*e.g*. phosphorus) in primary and detrital resources under nutrient enriched conditions should favour taxa with a high demand for this nutrient. With the nutrient demand of a taxon being correlated to the elemental composition of its body tissue (*e.g*. phosphorus content), such above described species shifts likely alter the overall community stoichiometry. However, studies addressing stoichiometry at community level are rare and most often restricted to lacustrine planktonic systems, single streams or limited experimental setups.

Relying on a stoichiometric database for >200 taxa and >1300 standardized sampling events of macroinvertebrate assemblages from the French national monitoring programme, we investigated the effect of water phosphorus and nitrogen load on stream macroinvertebrate community stoichiometry.

Community stoichiometry was significantly affected by water phosphorus concentration and the effect was strongest at low levels of nitrogen. While we could not confirm our hypothesis of increasing community %P (and decreasing C:P, N:P) with increasing water phosphorus concentrations for the overall community, it clearly followed this pattern for both Insecta and Malacostraca. General differences in the elemental composition among major taxonomic groups and a shift among these groups over the nutrient gradient probably explain the response of community stoichiometry. Our results show that assumptions from Ecological Stoichiometry Theory also hold at the community level, at least for two dominant taxa, and on a large spatial scale, with likely consequences for nutrient cycling and ecosystem function.

## Introduction

In a certain environment, taxa are selected based on their preferences (*e.g*. for food resources or substrates) and tolerances (*e.g*. towards pollution), thereby favouring those for which the mismatch between current environmental conditions and individual requirements is the lowest. Imbalances – for example between resource quality and organism nutrient demand – can occur under changing nutrient conditions (*e.g*. N, P) in the environment: while heterotrophs are quite homeostatic and keep a rather stable nutrient level, the elemental composition of basal resources changes accordingly and generally matches nutrient availabilities in the environment (Fink et al., 2006). For example, nutrient enrichment lowers C:nutrient ratios of primary producers, thus increasing their quality for herbivore consumers (Cross et al., 2003; Ventura et al., 2008). Similarly, detritus quality is increased at higher nutrient concentrations *via* increased microbial growth and nutrient storage leading to changed stoichiometry (Gulis & Suberkropp, 2003; Cross et al., 2007; Danger et al., 2016; Demi et al., 2018). For herbivore consumers, feeding on resources that largely deviate from the individual nutrient demand in any direction reduces the overall fitness (Boersma & Elser, 2006; Elser et al., 2016; Zhou & Declerck, 2019). Consequently, taxa whose nutrient demand is closer to the altered nutrient conditions should be favoured and nutrient concentrations in the environment should indirectly affect community structure. Nutrient-sensitive taxa that are adapted to low nutrient conditions are expected to disappear even at moderate nutrient increases, shifting the community towards more tolerant taxa (Ortiz & Puig, 2007; Friberg et al., 2010).

In previous studies, following increased phosphorus levels in the environment, abundances of phosphorus-rich taxa increased (Singer & Battin, 2007; Prater et al., 2015; Teurlincx et al., 2017), which increased the overall phosphorus content of macroinvertebrate and zooplankton communities (Singer & Battin, 2007; Teurlincx et al., 2017). Also over the long-term, community stoichiometry followed the decreasing phosphorus concentration in the environment with likely consequences for the nutrient cycling of a system (Beck et al., 2023). Body stoichiometry, that is the elemental composition, is assumed to be related to organisms nutrient demand (Sterner & Elser, 2002) and differs among taxonomic groups (Fagan et al., 2002; Evans-White et al., 2005; Teurlincx et al., 2017; Beck et al., 2022). With changing nutrient concentrations being a main stressor in worldwide ecosystems – either increasing due to anthropogenic activities (Falkowski et al., 2000; Peñuelas et al., 2013) or decreasing when counteracting management measures have been successful (Jeppesen et al., 2005; Floury et al., 2013; Latli et al., 2017), it seems indeed useful to benefit from the ultimate link towards nutrients that stoichiometric traits provide. Up to date, not many works have addressed stoichiometric questions considering consumer communities as a whole, and no study yet has explicitly investigated community stoichiometry on a large spatial scale. Considering its likely effect on nutrient storage over the long-term (Beck et al., 2023) and therewith higher trophic levels, it certainly seems worth investigating to close this research gab.

In this study, we aimed at analysing the effect of water phosphorus and nitrogen concentration on stream macroinvertebrate community stoichiometry. We considered coarse taxonomic groups (generally class) and functional feeding groups to investigate further details of these effects. We hypothesize that the concentrations of water nutrients will affect community stoichiometry. Specifically, we assume that community phosphorus content will be higher at sites where water phosphorus concentration is high, while community C:P and N:P stoichiometry decreases along this gradient. We expect to find this pattern for the stoichiometry of the complete community, as well as within each major macroinvertebrate group and it should be caused by a change in abundances among major taxonomic groups, and a within-group shift towards P-rich taxa, respectively. As overarching mechanism for the taxa shift, we assume altered quality and/or availability of basal resources under the different nutrient scenarios and a rather stable individual nutrient demand of consumers, represented by body stoichiometry (Beck et al., 2021). The community stoichiometry of primary consumers (detritivores, herbivores and omnivores) should be more strongly affected than that of secondary consumers (predators), whose resources are stoichiometrically more stable. Community stoichiometry within each feeding group should mainly follow the same pattern as described above for the complete community. We further integrate the level of water nitrogen into our analyses since in most systems not only absolute concentrations of both elements are changing but also their ratio (Peñuelas et al., 2013; Yan et al., 2016) and consequences on community composition have already been observed (Demi et al., 2019). Here, we expect higher impacts of phosphorus concentration under the highest nitrogen conditions, *i.e*, when invertebrate growth is likely limited by phosphorus.

We tested our hypotheses using information from a database containing stoichiometric information of >200 macroinvertebrate taxa and a dataset including >1300 site sampling events from a French national biomonitoring program, comprising macroinvertebrate abundance information and water chemistry. Macroinvertebrates are widely used in freshwater monitoring programs, thus providing extensive information on abundances and traits enabling such analyses.

## Material & Methods

### The data

Under the Water Framework Directive, benthic macroinvertebrate assemblages found in French wadeable streams are routinely monitored using standardized procedures (AFNOR 2016 for the field samplings; AFNOR 2020 for the sorting and identification steps). It allowed us to consolidate a large and standardized faunal dataset comprising the taxonomic composition and abundance of macroinvertebrate assemblages found in streams all over mainland France from 2004 to 2017, thus covering various regions and stream sizes (Figure 1). The general identification level was the genus level, whereas the family level was used for Diptera, Hirudinea and Turbellaria; Oligochaeta and Nematoda were identified as such.

**Figure 1:**
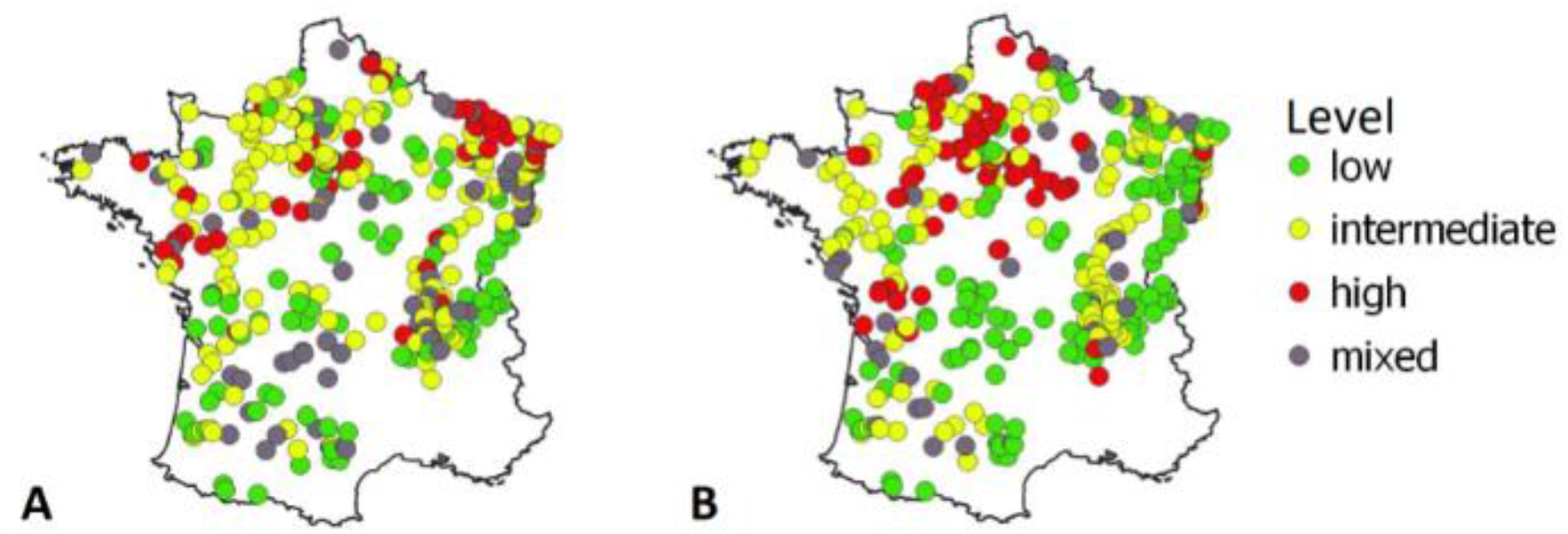
Geographic positions of sampling sites (n = 504) and their level regarding phosphate (A) and nitrogen (B) (“low”, “intermediate”, “high”) indicated by different colors. Grey dots (“mixed”) have been attributed when the nutrient level at a site changed between sampling events. Threshold concentrations for the displayed categories were the following: phosphorus: low: ≤ 0.05 mg/L, intermediate: > 0.05 & ≤ 0.2 mg/L, high > 0.2 mg/L; nitrogen: low: ≤ 10 mg/L, intermediate: > 10 & ≤ 25 mg/L, high: >25 mg/L.

The faunal dataset was complemented with a second dataset describing 17 pressure categories, including ten pressure categories linked to water quality (Mondy et al., 2012). Each pressure category was described using a set of five pressure levels, from worst to best: “bad”, “poor”, “moderate”, “good” and “high”. In order to avoid additional effects in heavily polluted sites, we first selected site sampling events (SSE; station x sampling date), which exhibited pressure levels either “high”, “good” or “moderate” for the following pressure categories: mineral micro-pollutants, pesticides, PAH, other organic micro-pollutants (*e.g*. chlorinated benzenes and phenols), suspended particulate matter, and nitrogen compounds (excluding nitrates). We then filtered by stream size, only keeping SSE from very small, small and intermediate streams (*i.e*. exhibiting a Strahler rank comprised between 1 and 4 (1 and 5 for sites located in the basin Loire-Bretagne)) for which data on the elemental composition of macroinvertebrate taxa was available (see *Data analysis*). Finally, we only kept SSE for which we had stoichiometric information available for at least 95% of the individuals. Based on these criteria, 1308 SSE were included in this study, with sampling years ranging between 2005 and 2017 and comprising 146 different taxa.

Last, we characterized the pressure level for phosphorus and nitrogen for our reduced dataset. Phosphorus level was used quantitatively and corresponds to the concentration of phosphates (PO_4_^3^-; mg/L). Nitrogen level refers to nitrate (NO_3_^-^ mg/L) concentration and was considered qualitatively as categorical variable representing three distinct water quality classes: low (**≤**10 mg/L), intermediate (10-25 mg/L) and high (>25 mg/L). It has to be noted that the quantitative values of phosphorus concentration contain a certain amount of imprecise values due to detection limits. In these cases, the true values might be lower than the reported value. Due to a lack of possibility to identify such values with certainty and to overcome the problem, we decided against excluding suspicious SSE and took the values as such. Most of these sites should be positioned at the real low end of the phosphorus concentration gradient, so that the overall trend we aim to find in the data should not be largely biased. Also, the nitrate values suffer from this problem, but due to the use of categories this does not pose a problem as all probably concerned SSE will fall into the same category (*i.e*. “low”, <10 mg/L). Distributions of water N and P concentrations are available in S1.

## Data analysis

### Community stoichiometry

Stoichiometric information, *i.e*. major nutrients (%C, %N, %P) and the corresponding molar ratios (C:N, C:P, N:P), of each taxon observed in our reduced dataset was obtained from a database (Beck *et al*., 2022). For each SSE, community stoichiometry was calculated as the abundance-weighted mean stoichiometry of taxa in the community. Thus, the relative abundance of each taxon (calculated as proportion of the sum of log[n+1] transformed abundances in the community) was multiplied by its corresponding stoichiometry value (%C, %N, %P, C:N, C:P, N:P) before summing up the values of all taxa in the community. The relative abundance thereby was based only on taxa for which the stoichiometric information was available, but covered ≥ 95% of the community in terms of individuals. Consequently, the number of taxa for a SSE can slightly vary between stoichiometric variables ranging from n= 134 (C:P and N:P) to n=147 (%P), but since we did not plan to compare stoichiometric variables among each other we decided for best data coverage. Values of the stoichiometric ratios (C:N, C:P, N:P) were log-transformed before the calculation to avoid biased results due to inert properties of ratio data as suggested by Isles (2020). Mass contents of the elements (%C, %N, %P) were used as such from the database.

Taxonomic group-wise and feeding group-wise community stoichiometric variables were calculated as described above, but by weighting the mean value of each taxon by its relative frequency within its major taxonomic or feeding group, rather than the complete community. The considered taxonomic groups were Bivalvia, Gastropoda, Hirudinea, Insecta, Malacostraca, Oligochaeta and Turbellaria. An updated (by the authors) version of the macroinvertebrate trait database from (Tachet et al., 2010) was used to assign each taxon to a feeding group, based on resource preference. This trait database provides fuzzy-coded trait information and contains the affinity (ranging from 0 = no affinity to 3 = high affinity) for each trait modality of a given trait. For a given taxon, affinities for each resource preference were first expressed as relative frequencies. We defined our feeding groups as follows, based on calculated relative frequencies of the affinities: (i) taxa with an affinity > 0.5 – i.e. the dominant share – for the resources “living micro-invertebrates” and “living macro-invertebrates” were considered predators; (ii) taxa with an affinity > 0.5 for plant-based resources (“living microphytes”, “living macrophytes”) were considered herbivores; (iii) taxa with an affinity > 0.5 for detritus-based resources (“dead plant”, “dead animal”, “fine detritus (*i.e*. size < 1 mm)”) were considered detritivores; and (iv) taxa with no affinity > 0.5 for any of these resource groups were considered omnivores. A taxalist including the feeding group classification is available in S10.

### Statistical analysis

Multiple linear models were separately applied for each stoichiometric variable to analyze the effect of water nutrient concentrations on community stoichiometry. Community stoichiometry was set as the dependent variable, whereas log(water P) and level of N were integrated as quantitative and categorical independent variables, respectively. An interaction term between the two independent variables was integrated and its contributing effect verified by comparison to a model without interaction term *via* the anova() function. If incorporating the interaction term did not improve the model, the simplest model was preferred. Model assumptions, which are normality of residuals and homoscedasticity, were checked graphically by inspecting QQ-plots.

The same models were applied to the sub-communities, setting one model per stoichiometric variable and taxonomic or feeding group. Due to the limited number of taxa in some taxonomic groups, only those groups that made the largest part of most communities and for which stoichiometric information was available for a sufficient number of taxa were considered for this part of the analysis (*i.e*. Gastropoda, Hirudinea, Insecta, Malacostraca). For the groups Gastropoda and Hirudinea, the assumption of normally distributed residuals was not met. Since this is most likely due to the limited number of taxa caught, which led to numerous identical values for group stoichiometry when only a few taxa of this group were present at a site, we still consider the results in this study under the corresponding reservation.

All analyses were conducted in R (version 3.5.3; R Core Team, 2020).

## Results

Please note that since our main hypotheses concerned phosphorus, we limit the detailed description to phosphorus-related stoichiometry and provide figures and results of %C, %N, and C:N for anyone interested in the supplementary material (Figures S2, S4, S6, Tables S3, S5, S7).

### Effect of water nutrient level on community stoichiometry

In all community stoichiometry values there was a significant effect of water P concentration and an interaction with the level of N (*i.e*. water P:level N), with the effect of water P being largest at low N-levels (Figure 2, Table S3).

**Figure 2:**
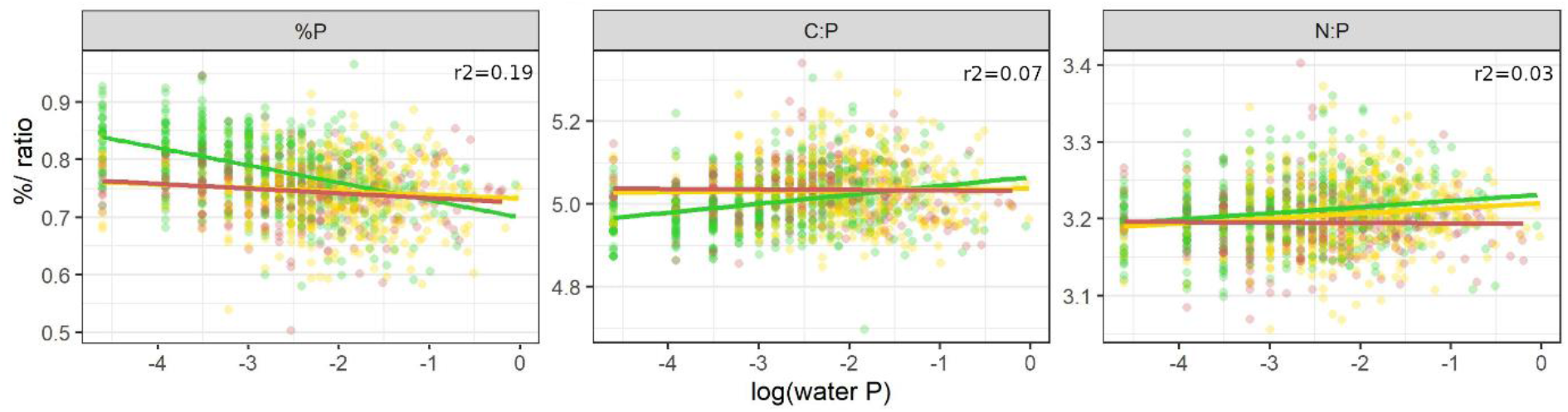
Community stoichiometry (%P, C:P, N:P) along a gradient of water P concentration (log scale). Regression lines allow different slopes and intercepts for different levels of water N. For each model, adjusted r2 values are provided and scattered points represent community stoichiometry for the different SSE with n = 519/ 501/ 288 SSE, respectively for low/ intermediate/ high N levels.Color codes for the level of water N: low (green), intermediate (yellow), high (red).

Community %P stoichiometry values significantly decreased with increasing water P, with the strength of the effect decreasing from low to intermediate/high N levels. In contrast, the values of stoichiometric ratios (C:P, N:P) significantly increased with increasing water P concentration at low and intermediate water N levels. Also here the effect was larger at low compared to intermediate levels of water N in both ratios. At high N levels however, the community ratios decreased but with a very low slope.

### Community stoichiometry within taxonomic groups

In all major macroinvertebrate taxonomic groups, community stoichiometry was significantly explained by water P concentration and/or nitrogen level.

Within Gastropoda, all community stoichiometry values were influenced by water phosphorus concentration with the effect depending on water N level and being strongest at intermediate N concentrations (Figure 3, Table S5). Community %P decreased with increasing water P concentration at intermediate N levels, but the responses under low and high N levels were insignificant. Community C:P and N:P were positively related to water P concentration at intermediate N, but tended to decrease under high N conditions. At low N levels, C:P ratios slightly increased while N:P ratios decreased along the water P gradient (both not significant).

**Figure 3:**
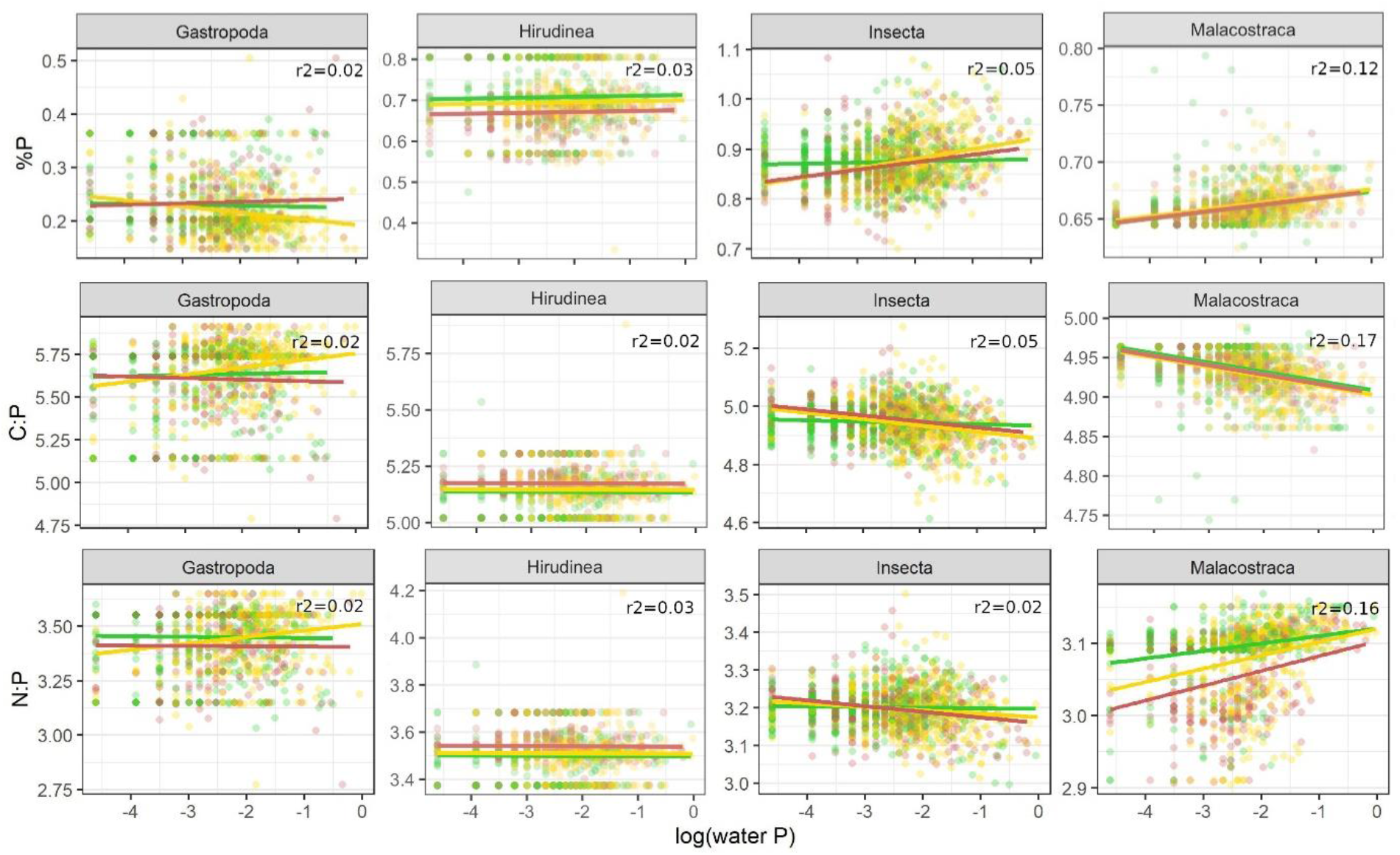
Community stoichiometry within taxonomic groups (%P, C:P, N:P) along water P concentration (log scale). Depending on the suggestion by model comparison, regression lines allow different slopes and intercepts for different levels of water N or assume the same slope for each level of water N. For each model, adjusted r2 values are provided; scattered data points represent community stoichiometry for the different SSE with n = 519/ 501/ 288 SSE, respectively for low/ intermediate/ high N levels. Color codes for the level of water N: low (green), intermediate (yellow), high (red).

Community stoichiometry of Hirudinea was not affected by water P concentration, but differed significantly between N levels. Community %P was lower under intermediate and high levels of N compared to low N conditions, while community C:P and N:P ratios showed the opposite pattern and values increased with N load.

Insect %P was positively related to water P concentration under all three levels of water N. The slope was low at low N sites but steepened at intermediate and high N sites. Stoichiometric ratios decreased with increasing water P concentration with the effect increasing with N load.

In Malacostraca, all stoichiometric variables were significantly affected by water P concentration but with an interaction between water nutrients only for community N:P. The positive effect on community N:P increased with increasing load of N. Also community %P was positively correlated to water P concentration, while the correlation of community C:P to water P concentration was negative with a slight but significant effet of water N load.

### Community stoichiometry within functional feeding groups

Within detritivores, all phosphorus-related stoichiometric variables significantly decreased with increasing water P concentration at low levels of N (Figure 4, Table S7). For %P, increasing levels of N decreased this effect, but it remained significant. Detritivore assemblage C:P values decreased with increasing water P and water N load. N:P ratios were positively correlated to water P concentration under intermediate and high N loads, but slightly declined under low N load.

**Figure 4:**
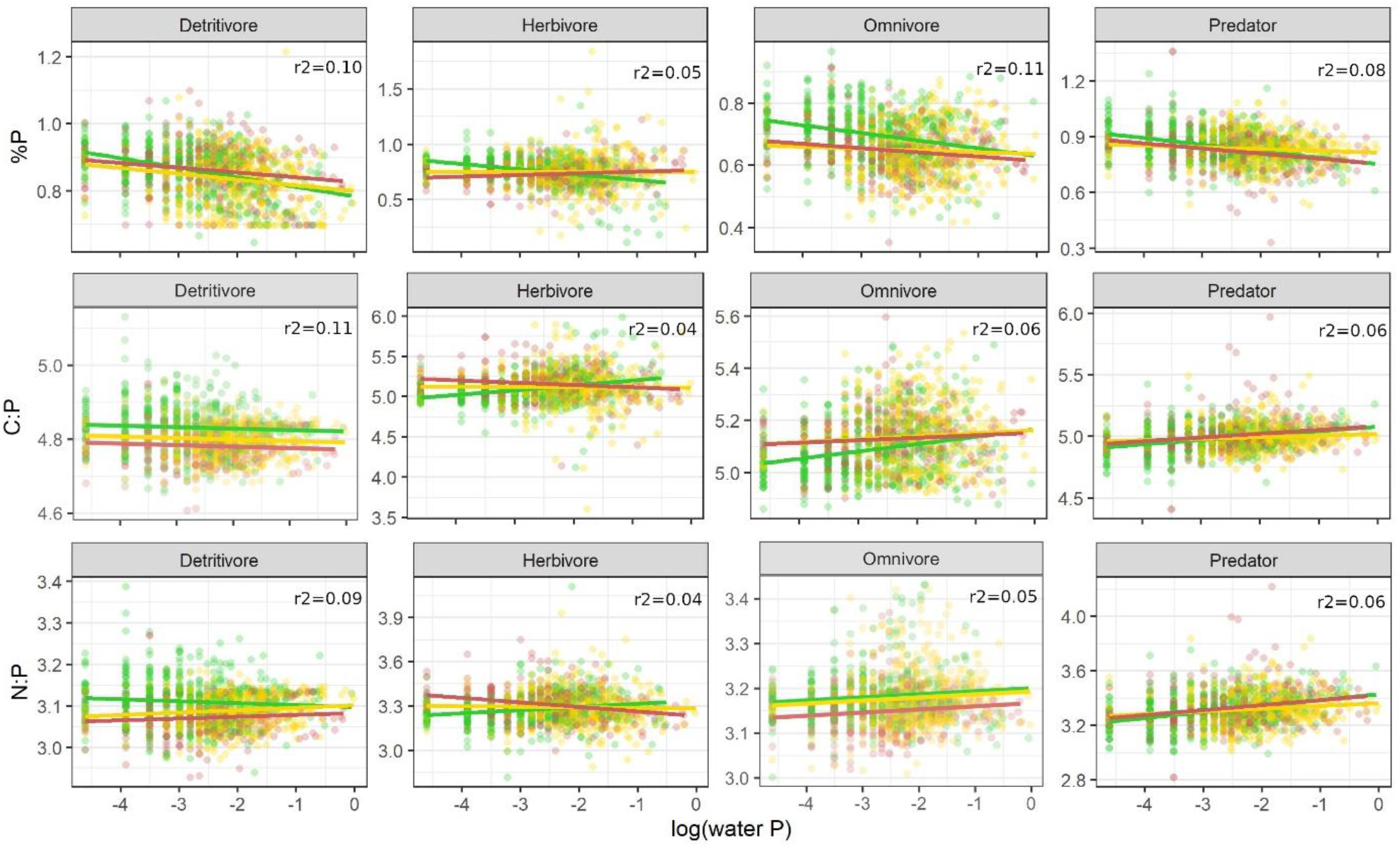
Community stoichiometry within functional feeding groups (%P, molar C:P, molar N:P) along water P concentration (log scale). Depending on the suggestion by model comparison, regression lines allow different slopes and intercepts for different levels of water N or assume the same slope for each level of water N. For each model, adjusted r2 values are provided; scattered data points represent community stoichiometry at the different SSE with n = 519/ 501/ 288 SSE, respectively for low/ intermediate/ high N levels. Color codes for the level of water N: low (green), intermediate (yellow), high (red).

In herbivores, increasing levels of water N opposed the effect of water P concentration on community stoichiometry values compared to low levels of N. %P was negatively correlated to water P concentration under low levels of N, while this correlation was positive under intermediate and high N load. Community C:P and N:P ratio both significantly increased with increasing water P under low N scenarios, but significantly decreased under higher levels of N.

In the omnivore assemblage %P significantly decreased and C:P ratio increased with increasing water P concentration. These effects were reduced under higher levels of N effect. N:P was significantly and positively correlated to water P concentration. There was no interaction with water N level, but under increased N load omnivore N:P was lower.

Predator assemblage %P significantly decreased with increasing water P concentration while C:P and N:P ratios significantly increased. For all three stoichiometry variables this effect was reduced under intermediate levels of water N.

### Proportions of taxonomic groups

Except for Turbellaria, the proportion of the taxonomic groups in the macroinvertebrate community depended on water phosphorus concentration and/or the level of nitrogen. While for Insecta, Malacostraca and Oligochaeta the effect of water P concentration on their proportion in the community depended on the level of water N, there was no such interaction in Bivalvia and Hirudinea. The proportion of Gastropoda was only affected by N load (Figure 5, Table S8).

**Figure 5:**
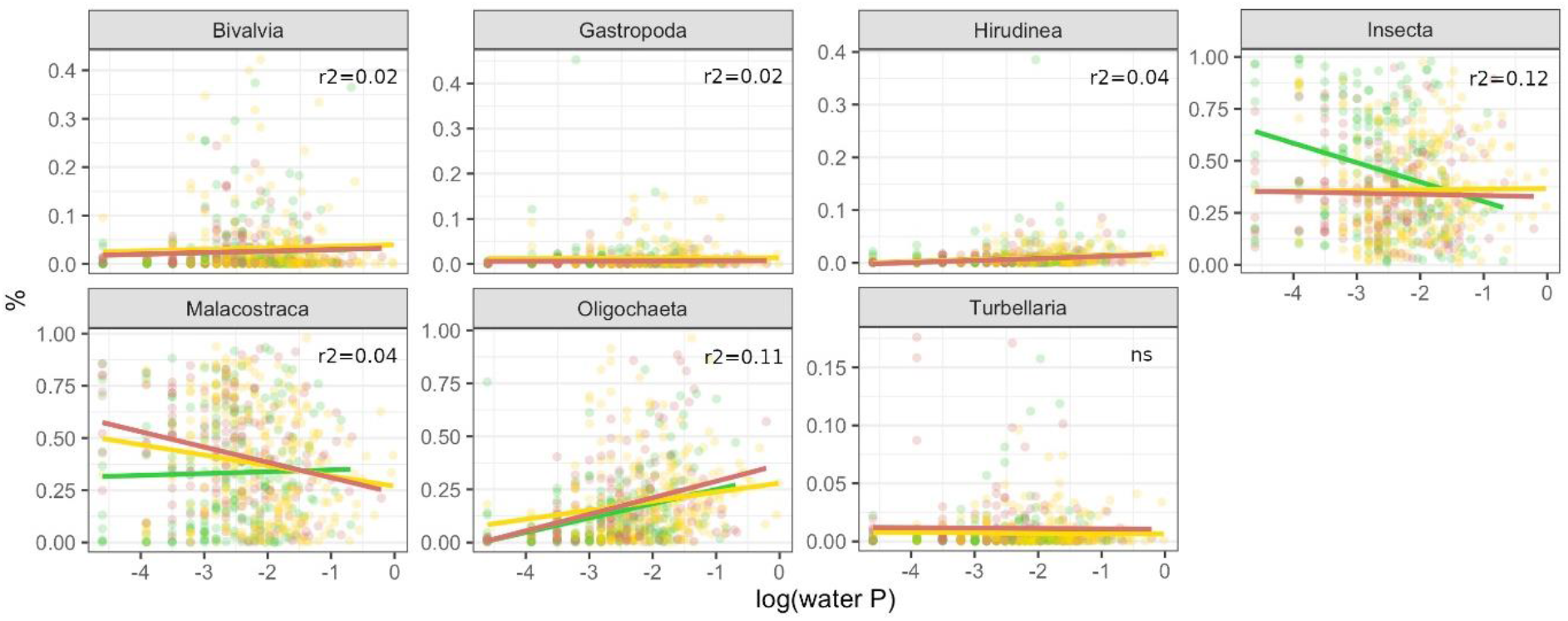
Proportion of macroinvertebrate groups within the macroinvertebrate community along water P concentration (log scale). Depending on the suggestion by model comparison, regression lines allow different slopes and intercepts for different levels of water N or assume the same slope for each level of water N. For each model, adjusted r2 values are provided; scattered data points represent community stoichiometry. Color codes for the level of water N: low (green), intermediate (yellow), high (red).

The proportion of Bivalvia and Hirudinea increased slightly but significantly with increasing water P concentration, and Bivalvia were additionally affected by the level of N. For Insecta, the effect of water P concentration was the largest when N levels were low. Their proportion then decreased significantly with increasing water P concentration, while it increased significantly at intermediate and at high N levels. Malacostraca proportion showed the opposite pattern: when N levels were low, the significant correlation between their proportion and water P concentration was positive, while it was negative when N levels were intermediate or high. The effect of water P then also was larger compared to low-N sites. The proportion of Oligochaeta significantly increased with increasing water P concentration, with the effect being largest at high N levels.

### Proportions of functional feeding groups

For all feeding groups, there was a significant interacting effect of water P concentration and the level of N on their relative abundance in the community (Figure 6, Table S8).

**Figure 6:**
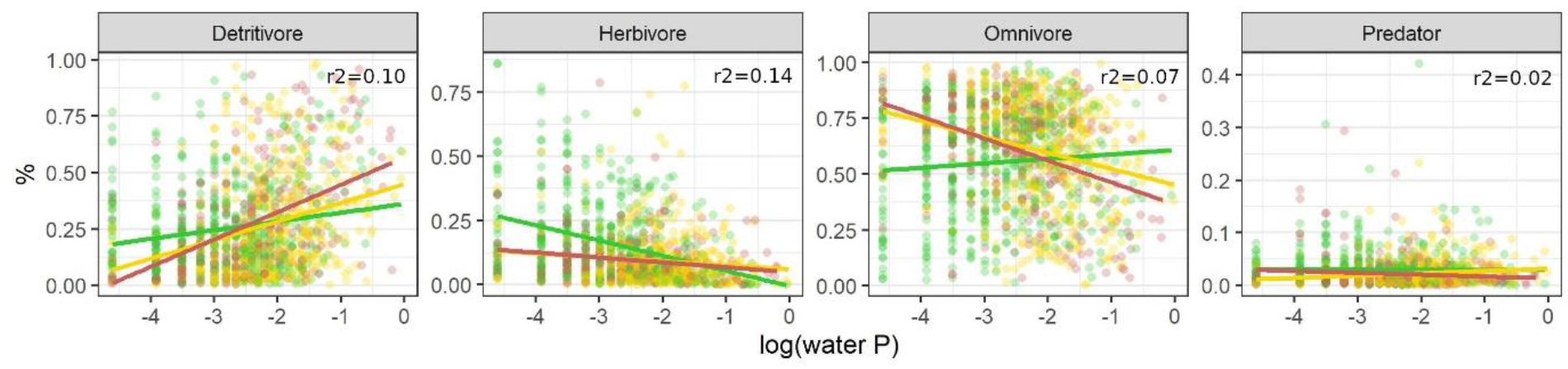
Proportion of functional feeding groups within the macroinvertebrate community along water P concentration (log scale). Depending on the suggestion by model comparison, regression lines allow different slopes and intercepts for different levels of water N or assume the same slope for each level of water N. For each model, adjusted r2 values provided; scattered data points represent community stoichiometry. Color codes for the level of water N: low (green), intermediate (yellow), high (red).

The proportion of detritivores increased significantly with increasing water P concentration at all the levels of N, but phosphorus had the largest effect at high levels of N. In contrast, herbivore proportion within the community significantly decreased over this gradient under all the water N scenarios, most strongly at low N levels. Omnivore proportion significantly decreased with increasing water P concentration under high and intermediate N levels, but significantly increased at low N levels. The relative abundance of predators was less affected by water nutrient conditions than the other groups. Its relationship with water P concentration was negative at low and high levels of N and positive at intermediate N levels.

## Discussion

Community stoichiometry of most taxonomic and functional groups was significantly affected by water P concentration and this effect depended on the level of N. Changes in taxonomic composition seem to be the main driver of this pattern, suggesting that species selection along a nutrient gradient also alters the nutrient pool provided by a consumer community.

Using stoichiometric trait information from a database assumes stoichiometric homeostasis in a way that intra-taxon variation is smaller than inter-taxon variation. Even though deviations from strict homeostasis with ontogeny (Back & King, 2013), body size (Naddafi et al., 2012; González et al., 2018) or stoichiometric differences between sexes (Back & King, 2013) have been reported for some taxa, the large number of data used in this analysis makes us confident that this assumption still leads to meaningful results. Compared to most works cited in this study that either conducted mesocosm experiments (Teurlincx et al., 2017), stream manipulations (Cross et al., 2003; Demi et al., 2019) or investigated a temporal nutrient gradient (Beck et al., 2023), we used data from large-scale field sampling of sites varying in nutrient levels. Rather than analysing the direct response of a community to a change in nutrient load, our approach thus addresses a broad spatial gradient and some general pattern. Although we reduced the risk of non-controlled confounding factors by selecting sites of similar sizes and excluding polluted streams, additional morphological or hydro-geographical stream characteristics might have reduced the stoichiometric trends in our analysis. Despite this noise, we have found a significant relationship in all stoichiometric variables, suggesting that there is a strong general link between environmental nutrient levels and community stoichiometry.

Interestingly, community stoichiometry values did not follow the P concentration in the water, which was in contrast to our first hypothesis that communities at high P sites should show higher %P and lower C:P and N:P than at low P sites. This is surprising, as two other studies also studying macroinvertebrates (Singer & Battin, 2007; Beck et al., 2023), found increasing community %P under high water P concentrations and similarly to Prater et al. (2015) reported shifts towards P-rich taxa. Community stoichiometry is most likely mainly driven by community composition and originates from the proportion of taxa that are rich and/or poor for a given nutrient. Teurlincx *et al*. (2017) demonstrated that a shift in plankton community C:P was caused by a change in community composition following nutrient enrichment, rather than stoichiometric plasticity of the individual taxa. Although there are stoichiometric variations among families or genera (McManamay et al., 2011; Morse et al., 2012; Mehler et al., 2013), those variations should be larger among classes or orders (Fagan et al., 2002; González et al., 2011; but see also González et al., 2018) due to general differences in body structure or metabolism between more distant lineages resulting in more contrasted nutrient requirements. In our study, the decrease in community %P with increasing water P went along with a sharp decrease of insect abundances at low N sites – a generally P-rich class with many species being sensitive to high nutrient levels (Friberg et al., 2010; Beck et al., 2022). Malacostraca and Oligochaeta – the most abundant taxonomic groups together with Insecta – are less rich in P and increased in proportion, which probably caused the overall decline in community %P stoichiometry. The proportion of Oligochaeta increased independently of N level (and especially at high N sites), which is not surprising as taxa of this group tend to dominate under high nutrient conditions (McCormick et al., 2004; Ortiz & Puig, 2007). Opposite trends in Malacostraca and Insecta – a moderate increase of Insecta while Malacostraca decline – could have reduced the effect on community %P under these conditions.

Also within macroinvertebrate groups community stoichiometry was altered, indicating that within-class taxonomic shifts also affect community stoichiometry. The responses however varied among groups.

For Insecta and Malacostraca we could confirm our hypothesis: there seems to be a shift towards P-rich taxa under high P concentrations, for example an increase in P-rich Diptera in Insecta while proportions of P-poor Coleoptera decreased (data not shown). Hirudinea seemed to be independent towards external nutrient drivers likely, at least partly, due to a hematophagous (parasitic) feeding strategy expressed widely within this group. The opposite responses of community stoichiometry under different levels of nitrogen as observed for Gastropoda, could indicate a biochemical co-limitation of nitrogen and phosphorus (Arrigo, 2005).

This effect of taxonomy on community stoichiometry seemed to over-shade that of feeding behaviour. Community stoichiometry values of feeding groups were less distinct from one another compared to taxonomic groups. They generally were affected in the same way by water nutrient concentrations and showed the same pattern as we found for the whole community. We could thus not confirm our hypothesis that community stoichiometry within primary consumers would respond more strongly to nutrient alterations than within secondary consumers. Evans-White et al. (2009) found mean C:P stoichiometry values decreasing under high water P conditions within some (shredders and collector-gatherers) – but not all – feeding groups. All this highlights that feeding groups are taxonomically diverse and, as several studies already demonstrated, taxonomy may better explain organism stoichiometry than its feeding group (Andrieux et al., 2020; Allgeier et al., 2020). Detritivores showed the strongest effect and were favoured under high P conditions. Herbivore proportions declined, suggesting a shift in functional composition following expectations based on their stoichiometric traits (González et al., 2011; McManamay et al., 2011). Compared to primary consumers, predators were only slightly affected by water nutrient conditions as it has been observed before (Evans-White et al., 2009). Although they can have a relatively high body phosphorus content (Cross et al., 2003; González et al., 2011), their prey are stoichiometrically more stable than the resources of primary consumers making prey identity probably more important. Some studies nevertheless reported increases in predators (Wimp et al., 2010; Murphy et al., 2012; Demi et al., 2019), probably as a consequence of increased prey availability.

The interaction with water N seems to be complicated, but in most cases the level of N did not affect the direction of the response in community stoichiometry towards increasing P concentrations, but the strength of this response. In contrast to our hypothesis, the community responded stronger under low levels of N, likely due to the strong and partly opposite responses in community composition, especially of the dominating groups Insecta and Malacostraca. Within Insecta, the effect of phosphorus concentration was stronger at higher levels of N, as expected, suggesting that under those conditions phosphorus was the limiting nutrient favoring P-rich taxa under high P condition. This response differed for Malacostraca where no interaction between N and P levels were evidenced. The rather strong response of Insecta stoichiometry to the N and P interaction could be related to the very high and variable N content of insects compared to other taxonomic groups, especially Malacostraca (Beck et al., 2022). Insecta could thus be more prone to N-limitation and stoichiometrically-induced taxonomic shifts in communities than other taxa, even at intermediate levels of N. In case of Hirudina, the hematophagous (parasitic) feeding strategy expressed widely within this group maybe makes them rather independant of such interaction and external nutrient drivers.

### Outlook – nutrient cycling and functioning

Ecological theory and some studies suggest a shift towards small-bodied taxa under high nutrient conditions (Singer & Battin, 2007; Demi et al., 2019), which could decrease the overall mass of phosphorus despite these taxa having higher body %P. A first estimation based on the data used in this study and relying to average body sizes, showed that although community %P decreased, the mass of phosphorus that is available in the community was not affected by water P or N load. One specificity of freshwater systems is the biomass loss due to emergence of semi-aquatic insects (Raitif et al., 2018). From the ecosystem perspective, this provides an energy and nutrient transfer between the aquatic and terrestrial systems, which can be substantial (Martin-Creuzburg et al., 2017; Bartrons et al., 2018; Raitif et al., 2018). For our data, the amount of phosphorus that should be lost from the system with emergence showed a strong decline when water phosphorus load increases (Figure S9), indicating that a larger part of phosphorus will remain in the system for a longer time. Accurately quantifying nutrient budgets could help understanding the effects of nutrient enrichment on nutrient cycling beyond community scale.

Due to (i) the link between body stoichiometry and biological traits (Beck et al., 2023), and (ii) taxa selection by water nutrient concentrations, one could assume a higher proportion of traits linked to high P body stoichiometry (*e.g*. fast growth rate, fast development, detritivory feeding behaviour) when the community phosphorus stoichiometry (%P) is high. Indeed, a long-term decline of phosphorus decreased overall community phosphorus content of macroinvertebrates and was associated to slow-developing taxa (Beck et al., 2023). Also decreases in taxon body size and life duration (Cross et al., 2003; Ortiz & Puig, 2007) were reported. This suggests a faster cycling of nutrients through this part of the assemblage compared to a system dominated by long-living taxa.

## Conclusion

We showed that taxonomic composition of stream macroinvertebrates was related to nutrient availability, leading to large effects on community stoichiometry. Even though the trends observed did not perfectly follow all of our assumptions, they demonstrate that assumptions from Ecological Stoichiometry Theory also hold at the community level and on a large spatial scale. Such changes mean an altered nutrient pool for higher trophic levels and were accompanied by shifts regarding feeding groups. Details of further consequences of such changes in community stoichiometry (*e.g*. functional trait composition) or that follow from this altered nutrient pool (*e.g*. regarding nutrient cycling) remain to be more deeply investigated.

## Acknowledgements

This research was supported by the Lorraine Université d’Excellence project “Stoichiometric Traits as Predictors of Community and Ecosystem response to global change” to M.D. and M.B. and by the StoichioMic ANR project (ANR-18-CE32-0003-01) to M.D. We thank everyone involved in the abiotic and biotic data collection of the WFD monitoring program, enabling us to use this great dataset.

## Data availability

Community and environmental data as well as the code used to run the analysis are available on zenodo (zenodo.org/records/10693755). Stoichiometric trait information are available in Beck et al. (2022).

