## Supplementary Material for "Effects of water nutrient concentrations on stream macroinvertebrate community stoichiometry: a large-scale study"

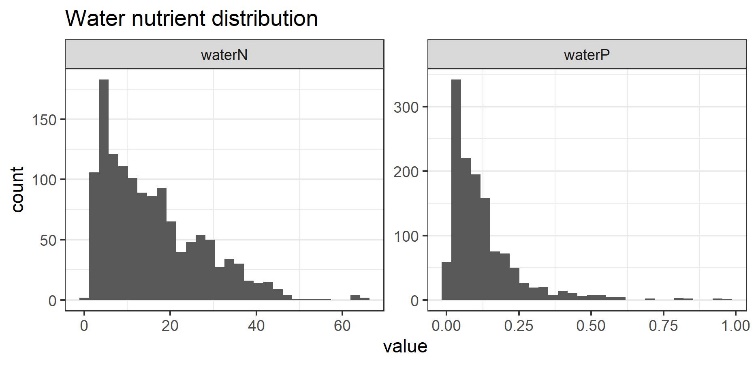

S1: Distribution of water nitrogen (NO_3_^-^ mg/L) and phosphorus (PO_4_^3-^) concentrations among study SSE.

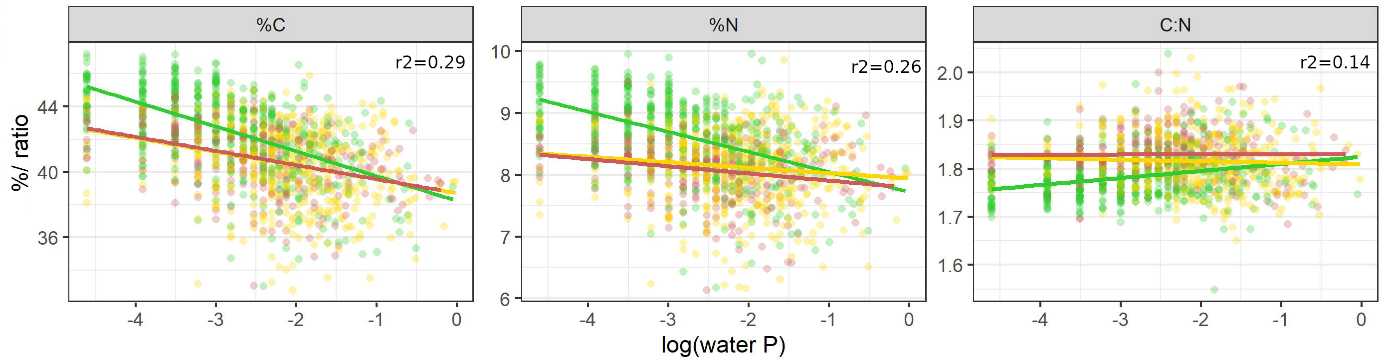

S2: Community stoichiometry (%C, %N, C:N) along water P concentration (log scale). Regression lines allow different slopes and intercept for different levels of water N. For each model, adjusted r2 values are provided; scattered points represent community stoichiometry with n=519/ 501/ 288 SSE, respectively for low/ intermediate/ high N level. Color codes for the level of water N: low (green), intermediate (yellow), high (red).

S3: Linear regression results for community stoichiometry. All models include an interaction term between water P concentration and level of N as suggested by model selection. For each model the slopes under the different N levels are given, as well as the adjusted model r2 and p-value. Asterisks indicate significance of effects with: * p < 0.05 ** p < 0.01 *** p < 0.001

|  | Effect | | | Slope | | |  |  |
| --- | --- | --- | --- | --- | --- | --- | --- | --- |
| Stoichio | P | N | NxP | low N | interm. N | high N | r2 | p value |
| %C | *** | *** | *** | -1.5127 | -0.8420 | -0.8605 | 0.2992 | 6.50E-99 |
| %N | *** | *** | *** | -0.3249 | -0.0878 | -0.1163 | 0.2552 | 8.04E-82 |
| %P | *** | *** | *** | -0.0303 | -0.0061 | -0.0082 | 0.1902 | 2.17E-58 |
| C:N | *** | *** | *** | 0.0146 | -0.0032 | 0.0002 | 0.1415 | 4.59E-42 |
| C:P | *** | *** | *** | 0.0215 | 0.0021 | -0.0012 | 0.0659 | 1.09E-18 |
| N:P | *** | *** | * | 0.0081 | 0.0066 | -0.0006 | 0.0317 | 5.94E-09 |

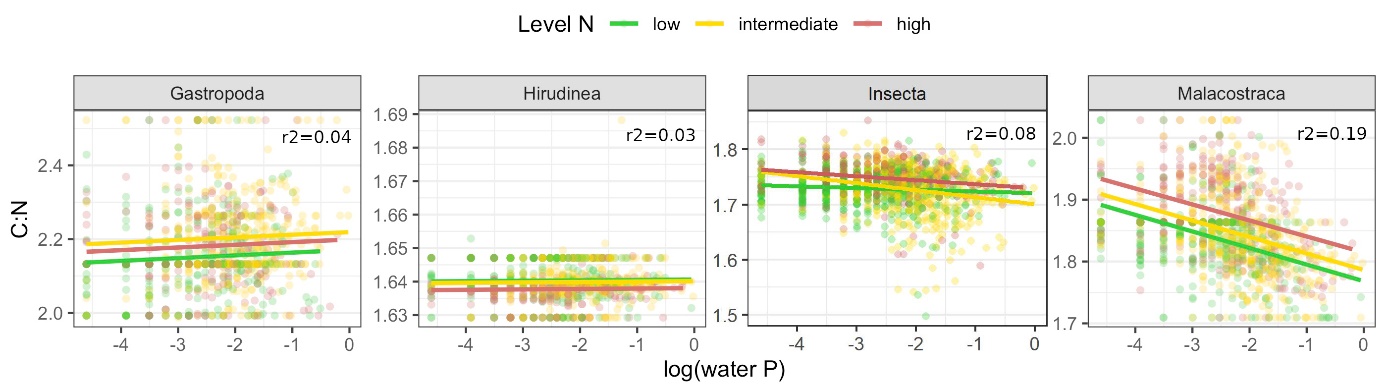

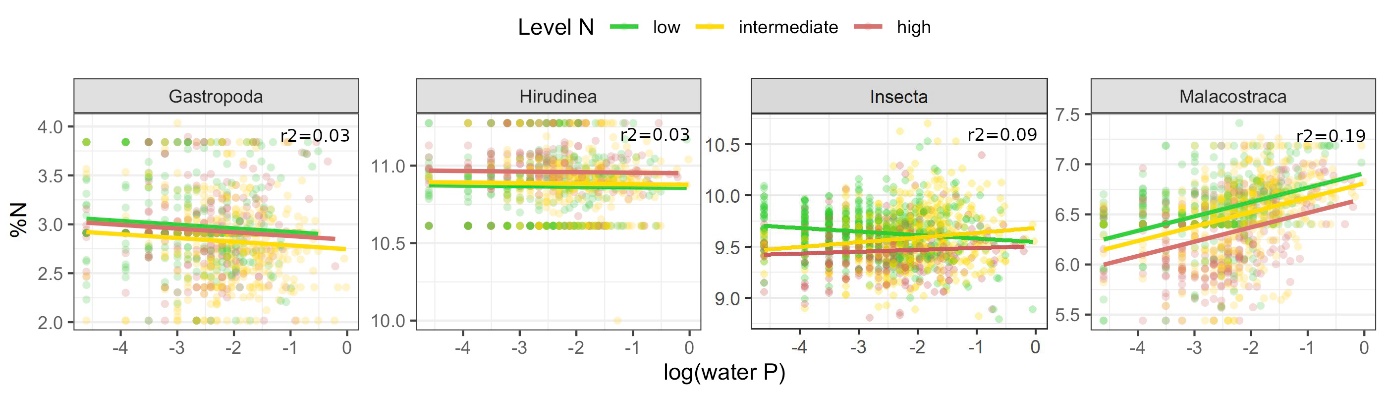

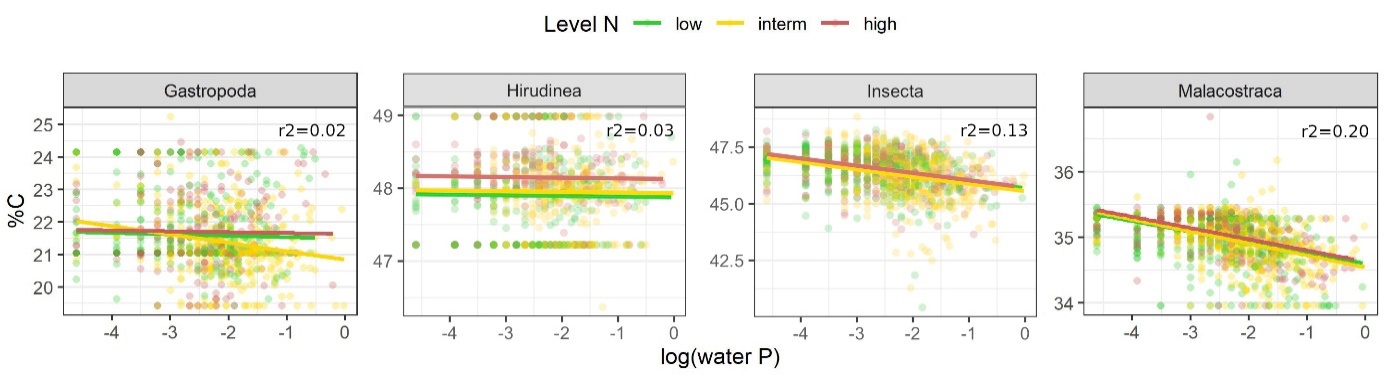

S4: Community stoichiometry within taxonomic groups (%C, %N, C:N) along water P concentration (log scale). Depending on the suggestion by model comparison, regression lines allow different slopes and intercepts for different levels of water N or assume the same slope for each level of water N. For each model, adjusted r2 values are provided; scattered data points represent community stoichiometry. Color codes for the level of water N: low (green), intermediate (yellow), high (red).

S5: Linear regression results for community stoichiometry within taxonomic groups. For each stoichiometric variable, the significance of effects (water P, water N, PxN interaction) and the slope under different water N loads are given, as well as the model adjusted r2 value and p-value. The choice of whether to include an interaction between water nutrients was made based on model comparison *via* anova() function. In models without interaction (“ns” for PxN), the slope indicated for “low N” refers to the general slope of the model. Asterisks indicate the significance with: * p < 0.05 ** p < 0.01 *** p < 0.001.

|  |  | Effect | | | Slope | | |  |  |
| --- | --- | --- | --- | --- | --- | --- | --- | --- | --- |
| Class | Stoichio | P | N | PxN | Low N | Interm. N | High N | r2 | p value |
| Gastropoda | %C | *** | ** | * | -0.0441 | -0.2568 | -0.0269 | 0.0227 | 2.02E-05 |
|  | %N | *** | *** | ns | -0.0380 |  |  | 0.0336 | 1.54E-08 |
|  | %P | ** | ** | * | -0.0019 | -0.0115 | 0.0028 | 0.0218 | 3.13E-05 |
|  | C:N | *** | *** | ns | 0.0079 |  |  | 0.0420 | 1.62E-10 |
|  | C:P | ** | ** | * | 0.0069 | 0.0416 | -0.0089 | 0.0211 | 4.27E-05 |
|  | N:P | ns | *** | * | -0.0027 | 0.0297 | -0.0014 | 0.0197 | 8.48E-05 |
| Hirudinea | %C | ns | *** | ns | -0.0091 |  |  | 0.0300 | 6.38E-07 |
|  | %N | ns | *** | ns | -0.0037 |  |  | 0.0299 | 6.68E-07 |
|  | %P | ns | *** | ns | 0.0034 |  |  | 0.0322 | 1.76E-07 |
|  | C:N | ns | *** | ns | 0.0001 |  |  | 0.0285 | 1.27E-06 |
|  | C:P | ns | *** | ns | -0.0004 |  |  | 0.0248 | 7.21E-06 |
|  | N:P | ns | *** | ns | -0.0006 |  |  | 0.0257 | 4.68E-06 |
| Insecta | %C | *** | * | ns | -0.3216 |  |  | 0.1278 | 4.93E-39 |
|  | %N | ns | *** | *** | -0.0342 | 0.0461 | 0.0182 | 0.0851 | 2.14E-24 |
|  | %P | *** | * | *** | 0.0022 | 0.0193 | 0.0150 | 0.0535 | 4.21E-15 |
|  | C:N | *** | *** | ** | -0.0031 | -0.0128 | -0.0072 | 0.0768 | 6.23E-22 |
|  | C:P | *** | ** | *** | -0.0045 | -0.0217 | -0.0207 | 0.0508 | 2.52E-14 |
|  | N:P | *** | ns | * | -0.0016 | -0.0091 | -0.0151 | 0.0185 | 2.03E-05 |
| Malacostraca | %C | *** | * | ns | -0.1713 |  |  | 0.2036 | 8.22E-63 |
|  | %N | *** | *** | ns | 0.1435 |  |  | 0.1864 | 5.96E-57 |
|  | %P | *** | ns | ns | 0.0060 |  |  | 0.1242 | 8.39E-37 |
|  | C:N | *** | *** | ns | -0.0269 |  |  | 0.1910 | 1.73E-58 |
|  | C:P | *** | ** | ns | -0.0119 |  |  | 0.1723 | 3.12E-52 |
|  | N:P | *** | *** | * | 0.0104 | 0.0185 | 0.0205 | 0.1565 | 1.31E-45 |

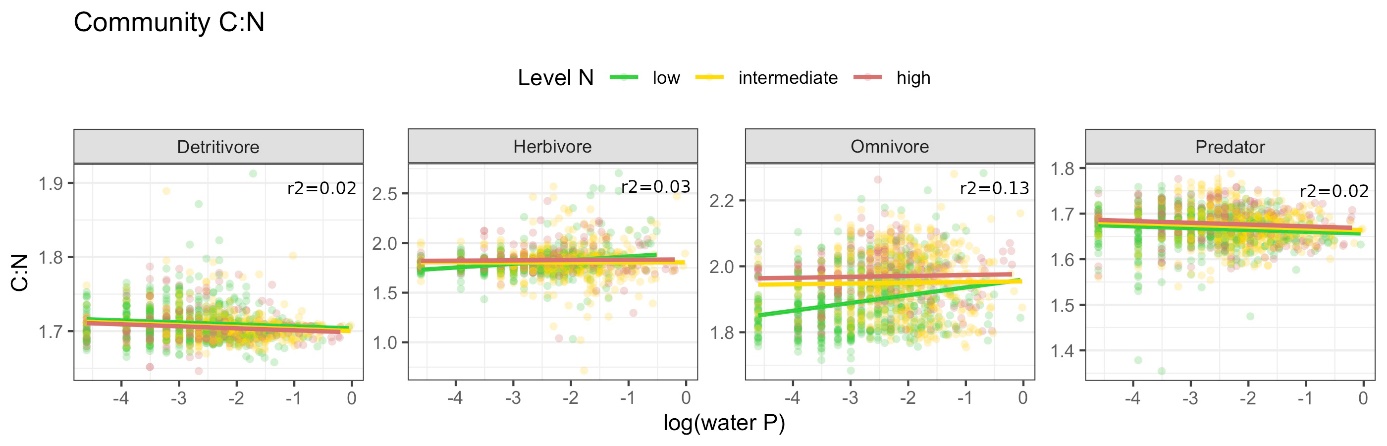

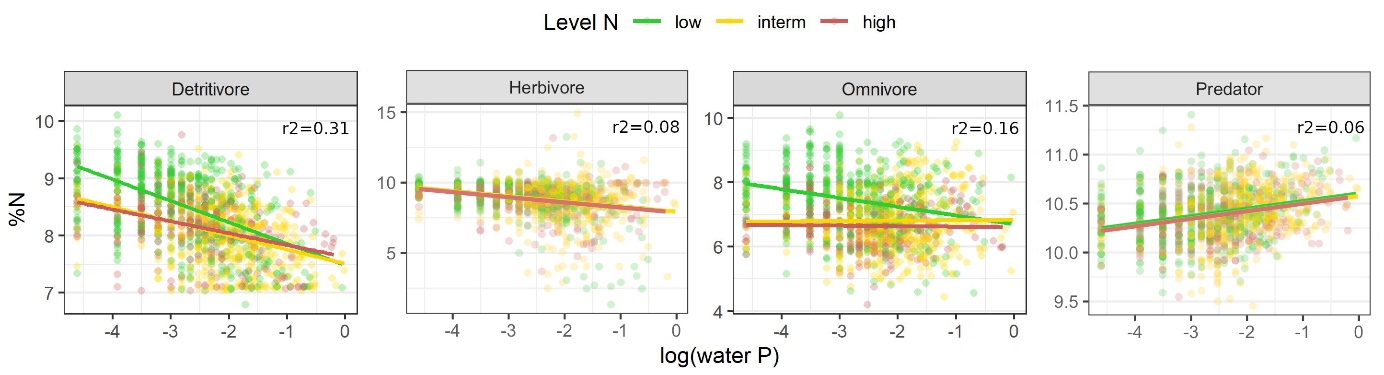

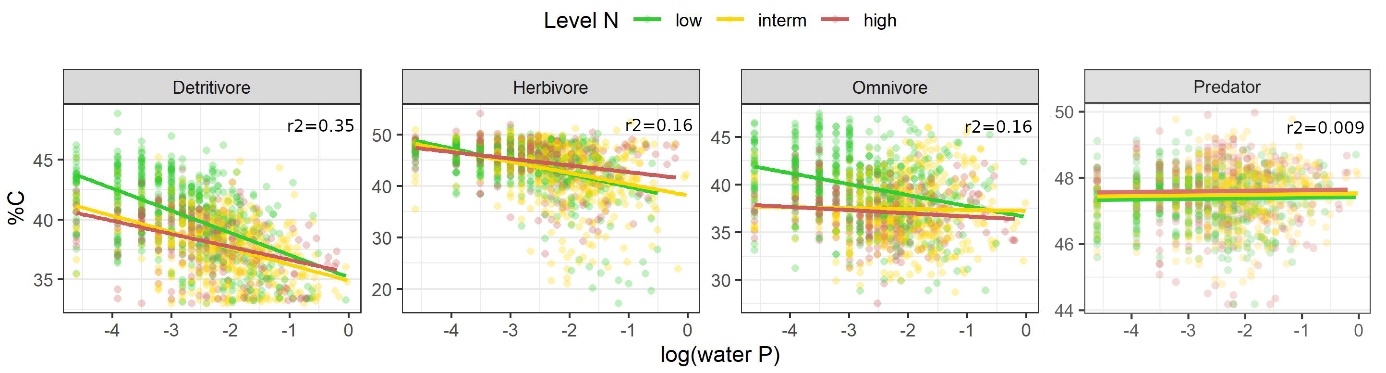

S6: Community stoichiometry (%C, %N, molar C:N) within functional feeding groups along water P concentration (log scale). Depending on the suggestion by model comparison, regression lines allow different slopes and intercepts for different levels of water N or assume the same slope for each level of water N. For each model, adjusted r2 values are provided; scattered data points represent community stoichiometry. Color codes for the level of water N: low (green), intermediate (yellow), high (red).

S7: Linear regression results for community stoichiometry within functional feeding groups. For each stoichiometric variable, the significance of effects (water P, water N, PxN interaction) and the slope under different water N loads are given. The choice of whether to include an interaction between water nutrients was made based on model comparison *via* anova() function. In models without interaction (“ns” for PxN), the slope indicated for “low N” refers to the general slope of the model. Asterisks indicate the significance of effects with: * p < 0.05 ** p < 0.01 *** p < 0.001.

|  |  | Effect | | | Slope | | |  |  |
| --- | --- | --- | --- | --- | --- | --- | --- | --- | --- |
|  | Stoichio | P | N | PxN | Low N | Interm. N | High N | r2 | p value |
| Detritivore | %C | *** | *** | *** | -1.8490 | -1.3574 | -1.0802 | 0.3456 | 3.40E-118 |
|  | %N | *** | *** | *** | -0.3747 | -0.2487 | -0.2074 | 0.3058 | 1.40E-101 |
|  | %P | *** | *** | * | -0.0286 | -0.0172 | -0.0142 | 0.0960 | 1.02E-27 |
|  | C:N | *** | ** | ns | -0.0026 |  |  | 0.0232 | 2.34E-07 |
|  | C:P | *** | *** | ns | -0.0077 |  |  | 0.1145 | 1.85E-33 |
|  | N:P | ns | *** | ** | -0.0046 | 0.0058 | 0.0044 | 0.0933 | 6.74E-27 |
| Herbivore | %C | *** | *** | ** | -2.4873 | -2.1683 | -1.3066 | 0.1579 | 5.73E-47 |
|  | %N | *** | ns | *** | -0.5587 | -0.2587 | -0.1568 | 0.0823 | 2.48E-23 |
|  | %P | *** | * | *** | -0.0482 | 0.0006 | 0.0150 | 0.0504 | 3.94E-14 |
|  | C:N | *** | * | *** | 0.0370 | -0.0002 | 0.0031 | 0.0252 | 3.94E-07 |
|  | C:P | *** | *** | *** | 0.0599 | -0.0035 | -0.0287 | 0.0442 | 2.57E-12 |
|  | N:P | ns | *** | *** | 0.0213 | -0.0039 | -0.0306 | 0.0364 | 3.75E-10 |
| Omnivore | %C | *** | *** | *** | -1.1524 | -0.0957 | -0.3325 | 0.1583 | 1.43E-47 |
|  | %N | *** | *** | *** | -0.2732 | 0.0129 | -0.0165 | 0.1641 | 1.65E-49 |
|  | %P | *** | *** | ** | -0.0247 | -0.0059 | -0.0138 | 0.1126 | 7.47E-33 |
|  | C:N | *** | *** | *** | 0.0236 | 0.0022 | 0.0028 | 0.1317 | 6.79E-39 |
|  | C:P | *** | *** | * | 0.0282 | 0.0114 | 0.0096 | 0.0604 | 4.48E-17 |
|  | N:P | ** | *** | ns | 0.0067 |  |  | 0.0501 | 4.38E-15 |
| Predator | %C | ns | ** | ns | 0.0184 |  |  | 0.0094 | 1.61 E-3 |
|  | %N | *** | ns | ns | 0.0772 |  |  | 0.0643 | 2.82E-19 |
|  | %P | *** | ** | *** | -0.0353 | -0.0094 | -0.0276 | 0.0835 | 7.28E-24 |
|  | C:N | * | *** | ns | -0.0037 |  |  | 0.0151 | 4.40E-05 |
|  | C:P | *** | * | * | 0.0360 | 0.0127 | 0.0308 | 0.0554 | 1.35E-15 |
|  | N:P | *** | ns | * | 0.0437 | 0.0200 | 0.0366 | 0.0639 | 4.70E-18 |

S8: Linear regression results for the proportion of taxonomic and feeding groups within the macroinvertebrate community. For each group, the significance of effects (water P, water N, PxN interaction) and the slope under different water N loads are given, as well as the model adjusted r2 value and p-value. The choice of whether to include an interaction between water nutrients was made based on model comparison *via* anova() function. In models without interaction (“ns” for PxN), the slope indicated for “low N” refers to the general slope of the model. Asterisks indicate the significance of effects with: * p < 0.05 ** p < 0.01 *** p < 0.001.

|  | Effect | | | Slope | | |  |  |
| --- | --- | --- | --- | --- | --- | --- | --- | --- |
| Group | P | N | PxN | Low N | Interm. N | High N | R2 | p.value |
| Bivalvia | * | * | ns | 0.0081 |  |  | 0.0153 | 2. 28E-03 |
| Gastropoda | ns | ** | ns | 0.0021 |  |  | 0.0180 | 8. 05E-04 |
| Hirudinea | *** | ns | ns | 0.0044 |  |  | 0.0367 | 9.28E-07 |
| Insecta | *** | *** | *** | -0.1038 | 0.0305 | 0.0085 | 0.1171 | 2.73E-41 |
| Malacostraca | *** | *** | *** | 0.0293 | -0.0719 | -0.1128 | 0.0380 | 9.97E-18 |
| Oligochaeta | *** | ** | * | 0.0533 | 0.0432 | 0.0983 | 0.1117 | 1.23E-30 |
| Turbellaria (ns) | ns | ns | ns | -0.0002 |  |  | -0.0008 | 0.49 |
| Detritivore | *** | ns | *** | 0.0387 | 0.0824 | 0.1212 | 0.1047 | 2.03E-30 |
| Herbivore | *** | *** | *** | -0.0590 | -0.0157 | -0.0190 | 0.1405 | 9.01E-42 |
| Omnivore | *** | *** | *** | 0.0198 | -0.0715 | -0.0983 | 0.0677 | 2.97E-19 |
| Predator | ns | *** | ** | -0.0003 | 0.0044 | -0.0035 | 0.0191 | 1.42E-05 |

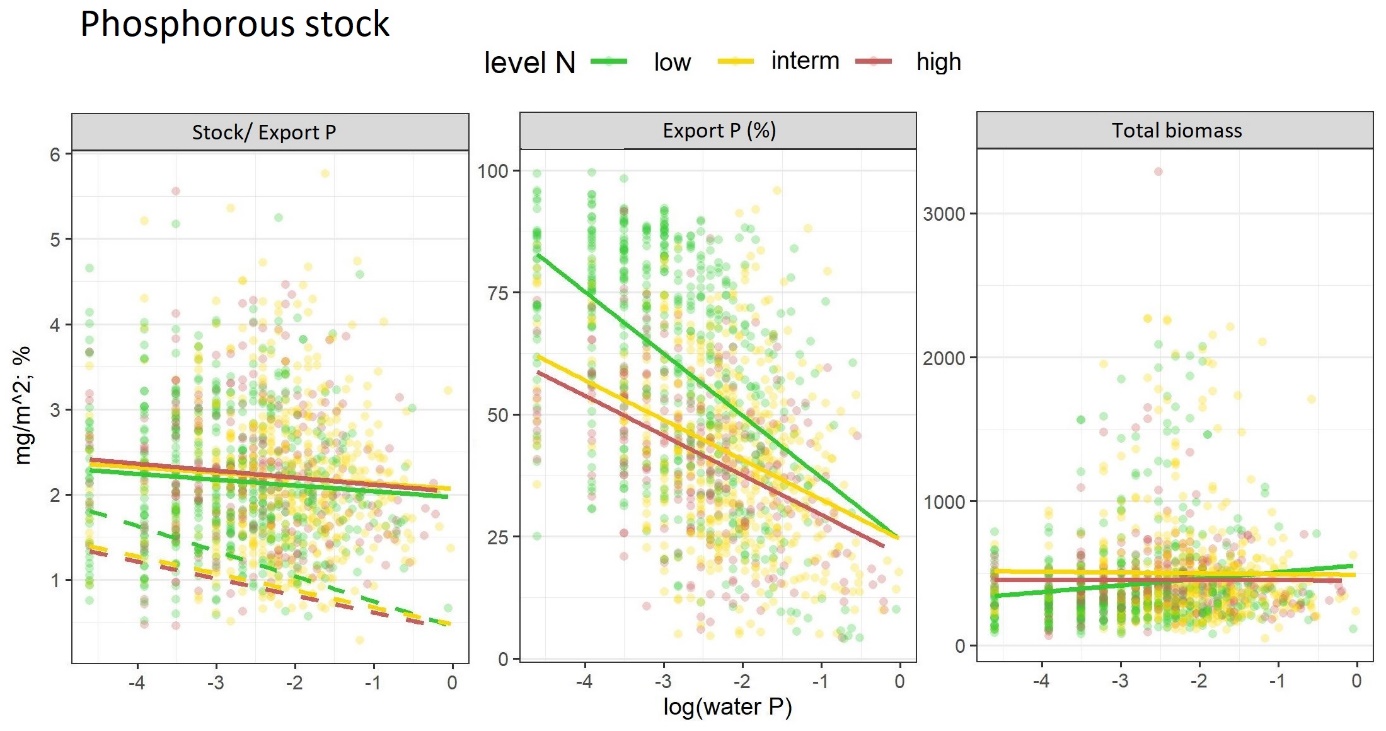

S9: Estimations of phosphorus stock (mg/m^2^), export (% of stored P), and total biomass (mg/m^2^) along water P concentration (log scale). Total biomass was calculated by multiplying each taxon’s abundance by its mean taxon biomass (unpublished database that is based on a literature research and several sampling campaigns). Phosphorus stock (solid lines in first panel) was calculated by multiplying biomass weighted abundances by the taxon’s phosphorus content, The phosphorus stock considering only taxa that will emerge into terrestrial adults gives the phosphorus export (dashed line in first panel). The %P of this export is obtained by calculating the group stoichiometry of those taxa. Scattered data points represent raw data (calculated community stoichiometry at the different sites). Color codes for the level of water N: low (green), intermediate (yellow), high (red).

S10: List of taxa included in the analysis, i.e. for which stoichiometric information was available. For each taxon, the assumed feeding group and taxonomy (class, order, family, genus) is provided.

| Taxon | feeding_group | class | order | family | genus |
| --- | --- | --- | --- | --- | --- |
| Hydracarina | Omnivore | Arachnida | Hydracarina | NA | NA |
| *Corbicula* | Omnivore | Bivalvia | NA | Corbiculidae | *Corbicula* |
| *Dreissena* | Omnivore | Bivalvia | NA | Dreissenidae | *Dreissena* |
| Dreissenidae | Omnivore | Bivalvia | NA | Dreissenidae | NA |
| *Pisidium* | Omnivore | Bivalvia | NA | Sphaeriidae | *Pisidium* |
| Sphaeriidae | Herbivore | Bivalvia | NA | Sphaeriidae | NA |
| *Ancylus* | Herbivore | Gastropoda | NA | Ancylidae | *Ancylus* |
| *Bithynia* | Omnivore | Gastropoda | NA | Bithyniidae | *Bithynia* |
| *Galba* | Herbivore | Gastropoda | NA | Lymnaeidae | *Galba* |
| *Radix* | Omnivore | Gastropoda | NA | Lymnaeidae | *Radix* |
| *Stagnicola* | Herbivore | Gastropoda | NA | Lymnaeidae | *Stagnicola* |
| Lymnaeidae | Omnivore | Gastropoda | NA | Lymnaeidae | NA |
| Physidae | Herbivore | Gastropoda | NA | Physidae | NA |
| Planorbidae | Omnivore | Gastropoda | NA | Planorbidae | NA |
| Hirudidae | Predator | Hirudinea | Gnathobdelliformes | Hirudidae | NA |
| Erpobdellidae | Predator | Hirudinea | Pharyngobdelliformes | Erpobdellidae | NA |
| Glossiphoniidae | Predator | Hirudinea | Rhynchobdelliformes | Glossiphoniidae | NA |
| Piscicolidae | Predator | Hirudinea | Rhynchobdelliformes | Piscicolidae | NA |
| Hydroporinae | Predator | Insecta | Coleoptera | Dytiscidae | NA |
| Laccophilinae | Predator | Insecta | Coleoptera | Dytiscidae | NA |
| Colymbetinae | Predator | Insecta | Coleoptera | Dytiscidae | NA |
| Dytiscidae | Predator | Insecta | Coleoptera | Dytiscidae | NA |
| *Elmis* | Herbivore | Insecta | Coleoptera | Elmidae | *Elmis* |
| *Limnius* | Herbivore | Insecta | Coleoptera | Elmidae | *Limnius* |
| *Macronychus* | Herbivore | Insecta | Coleoptera | Elmidae | *Macronychus* |
| *Oulimnius* | Herbivore | Insecta | Coleoptera | Elmidae | *Oulimnius* |
| *Stenelmis* | Herbivore | Insecta | Coleoptera | Elmidae | *Stenelmis* |
| Elmidae | Herbivore | Insecta | Coleoptera | Elmidae | NA |
| *Haliplus* | Herbivore | Insecta | Coleoptera | Haliplidae | *Haliplus* |
| Haliplidae | Herbivore | Insecta | Coleoptera | Haliplidae | NA |
| Hydraenidae | Herbivore | Insecta | Coleoptera | Hydraenidae | NA |
| Hydrophilinae | Predator | Insecta | Coleoptera | Hydrophilidae | NA |
| Hydrophilidae | Predator | Insecta | Coleoptera | Hydrophilidae | NA |
| *Noterus* | Predator | Insecta | Coleoptera | Noteridae | *Noterus* |
| Scirtidae | Herbivore | Insecta | Coleoptera | Scirtidae | NA |
| Athericidae | Predator | Insecta | Diptera | Athericidae | NA |
| Ceratopogonidae | Herbivore | Insecta | Diptera | Ceratopogonidae | NA |
| Chironomidae | Omnivore | Insecta | Diptera | Chironomidae | NA |
| Dixidae | Omnivore | Insecta | Diptera | Dixidae | NA |
| Empididae | Predator | Insecta | Diptera | Empididae | NA |
| Limoniidae | Detritivore | Insecta | Diptera | Limoniidae | NA |
| Psychodidae | Detritivore | Insecta | Diptera | Psychodidae | NA |
| Ptychopteridae | Detritivore | Insecta | Diptera | Ptychopteridae | NA |
| Simuliidae | Detritivore | Insecta | Diptera | Simuliidae | NA |
| Stratiomyidae | Omnivore | Insecta | Diptera | Stratiomyidae | NA |
| Tabanidae | Predator | Insecta | Diptera | Tabanidae | NA |
| Tipulidae | Omnivore | Insecta | Diptera | Tipulidae | NA |
| *Baetis* | Herbivore | Insecta | Ephemeroptera | Baetidae | *Baetis* |
| Baetidae | Omnivore | Insecta | Ephemeroptera | Baetidae | NA |
| *Caenis* | Detritivore | Insecta | Ephemeroptera | Caenidae | *Caenis* |
| Caenidae | Detritivore | Insecta | Ephemeroptera | Caenidae | NA |
| *Ephemerella* | Herbivore | Insecta | Ephemeroptera | Ephemerellidae | *Ephemerella* |
| *Torleya* | Detritivore | Insecta | Ephemeroptera | Ephemerellidae | *Torleya* |
| Ephemerellidae | Detritivore | Insecta | Ephemeroptera | Ephemerellidae | NA |
| *Ephemera* | Detritivore | Insecta | Ephemeroptera | Ephemeridae | *Ephemera* |
| *Ecdyonurus* | Detritivore | Insecta | Ephemeroptera | Heptageniidae | *Ecdyonurus* |
| *Epeorus* | Herbivore | Insecta | Ephemeroptera | Heptageniidae | *Epeorus* |
| *Heptagenia* | Detritivore | Insecta | Ephemeroptera | Heptageniidae | *Heptagenia* |
| *Rhithrogena* | Herbivore | Insecta | Ephemeroptera | Heptageniidae | *Rhithrogena* |
| Heptageniidae | Herbivore | Insecta | Ephemeroptera | Heptageniidae | NA |
| *Habroleptoides* | Detritivore | Insecta | Ephemeroptera | Leptophlebiidae | *Habroleptoides* |
| *Habrophlebia* | Detritivore | Insecta | Ephemeroptera | Leptophlebiidae | *Habrophlebia* |
| *Leptophlebia* | Detritivore | Insecta | Ephemeroptera | Leptophlebiidae | *Leptophlebia* |
| *Paraleptophlebia* | Detritivore | Insecta | Ephemeroptera | Leptophlebiidae | *Paraleptophlebia* |
| Leptophlebiidae | Detritivore | Insecta | Ephemeroptera | Leptophlebiidae | NA |
| *Siphlonurus* | Omnivore | Insecta | Ephemeroptera | Siphlonuridae | *Siphlonurus* |
| *Aphelocheirus* | Predator | Insecta | Hemiptera | Aphelocheiridae | *Aphelocheirus* |
| Nepidae | Predator | Insecta | Hemiptera | Nepidae | NA |
| Veliidae | Predator | Insecta | Hemiptera | Veliidae | NA |
| *Sialis* | Predator | Insecta | Megaloptera | Sialidae | *Sialis* |
| *Aeshna* | Predator | Insecta | Odonata | Aeshnidae | *Aeshna* |
| *Boyeria* | Predator | Insecta | Odonata | Aeshnidae | *Boyeria* |
| Aeshnidae | Predator | Insecta | Odonata | Aeshnidae | NA |
| *Calopteryx* | Predator | Insecta | Odonata | Calopterygidae | *Calopteryx* |
| Coenagrionidae | Predator | Insecta | Odonata | Coenagrionidae | NA |
| *Cordulegaster* | Predator | Insecta | Odonata | Cordulegasteridae | *Cordulegaster* |
| *Gomphus* | Predator | Insecta | Odonata | Gomphidae | *Gomphus* |
| *Onychogomphus* | Predator | Insecta | Odonata | Gomphidae | *Onychogomphus* |
| Gomphidae | Predator | Insecta | Odonata | Gomphidae | NA |
| *Libellula* | Predator | Insecta | Odonata | Libellulidae | *Libellula* |
| Libellulidae | Predator | Insecta | Odonata | Libellulidae | NA |
| *Platycnemis* | Predator | Insecta | Odonata | Platycnemididae | *Platycnemis* |
| *Capnia* | Herbivore | Insecta | Plecoptera | Capniidae | *Capnia* |
| Capniidae | Detritivore | Insecta | Plecoptera | Capniidae | NA |
| *Siphonoperla* | Predator | Insecta | Plecoptera | Chloroperlidae | *Siphonoperla* |
| Chloroperlidae | Predator | Insecta | Plecoptera | Chloroperlidae | NA |
| *Leuctra* | Herbivore | Insecta | Plecoptera | Leuctridae | *Leuctra* |
| Leuctridae | Omnivore | Insecta | Plecoptera | Leuctridae | NA |
| *Amphinemura* | Detritivore | Insecta | Plecoptera | Nemouridae | *Amphinemura* |
| *Nemoura* | Detritivore | Insecta | Plecoptera | Nemouridae | *Nemoura* |
| *Protonemura* | Detritivore | Insecta | Plecoptera | Nemouridae | *Protonemura* |
| Nemouridae | Detritivore | Insecta | Plecoptera | Nemouridae | NA |
| *Dinocras* | Predator | Insecta | Plecoptera | Perlidae | *Dinocras* |
| *Perla* | Predator | Insecta | Plecoptera | Perlidae | *Perla* |
| Perlidae | Predator | Insecta | Plecoptera | Perlidae | NA |
| *Isoperla* | Predator | Insecta | Plecoptera | Perlodidae | *Isoperla* |
| *Perlodes* | Predator | Insecta | Plecoptera | Perlodidae | *Perlodes* |
| Perlodidae | Predator | Insecta | Plecoptera | Perlodidae | NA |
| *Brachyptera* | Herbivore | Insecta | Plecoptera | Taeniopterygidae | *Brachyptera* |
| *Taeniopteryx* | Detritivore | Insecta | Plecoptera | Taeniopterygidae | *Taeniopteryx* |
| Taeniopterygidae | Detritivore | Insecta | Plecoptera | Taeniopterygidae | NA |
| *Brachycentrus* | Herbivore | Insecta | Trichoptera | Brachycentridae | *Brachycentrus* |
| Brachycentridae | Herbivore | Insecta | Trichoptera | Brachycentridae | NA |
| *Agapetus* | Herbivore | Insecta | Trichoptera | Glossosomatidae | *Agapetus* |
| *Glossosoma* | Herbivore | Insecta | Trichoptera | Glossosomatidae | *Glossosoma* |
| Glossosomatidae | Herbivore | Insecta | Trichoptera | Glossosomatidae | NA |
| *Goera* | Omnivore | Insecta | Trichoptera | Goeridae | *Goera* |
| Goeridae | Herbivore | Insecta | Trichoptera | Goeridae | NA |
| *Diplectrona* | Omnivore | Insecta | Trichoptera | Hydropsychidae | *Diplectrona* |
| *Hydropsyche* | Omnivore | Insecta | Trichoptera | Hydropsychidae | *Hydropsyche* |
| Hydropsychidae | Omnivore | Insecta | Trichoptera | Hydropsychidae | NA |
| *Ptilocolepus* | Herbivore | Insecta | Trichoptera | Hydroptilidae | *Ptilocolepus* |
| Hydroptilidae | Herbivore | Insecta | Trichoptera | Hydroptilidae | NA |
| *Lepidostoma* | Detritivore | Insecta | Trichoptera | Lepidostomatidae | *Lepidostoma* |
| Lepidostomatidae | Detritivore | Insecta | Trichoptera | Lepidostomatidae | NA |
| *Adicella* | Herbivore | Insecta | Trichoptera | Leptoceridae | *Adicella* |
| *Athripsodes* | Herbivore | Insecta | Trichoptera | Leptoceridae | *Athripsodes* |
| *Mystacides* | Herbivore | Insecta | Trichoptera | Leptoceridae | *Mystacides* |
| *Oecetis* | Herbivore | Insecta | Trichoptera | Leptoceridae | *Oecetis* |
| Leptoceridae | Herbivore | Insecta | Trichoptera | Leptoceridae | NA |
| Limnephilidae | Herbivore | Insecta | Trichoptera | Limnephilidae | NA |
| Drusinae | Herbivore | Insecta | Trichoptera | Limnephilidae | NA |
| Limnephilinae | Herbivore | Insecta | Trichoptera | Limnephilidae | NA |
| *Odontocerum* | Predator | Insecta | Trichoptera | Odontoceridae | *Odontocerum* |
| *Chimarra* | Omnivore | Insecta | Trichoptera | Philopotamidae | *Chimarra* |
| *Philopotamus* | Herbivore | Insecta | Trichoptera | Philopotamidae | *Philopotamus* |
| Philopotamidae | Herbivore | Insecta | Trichoptera | Philopotamidae | NA |
| *Plectrocnemia* | Predator | Insecta | Trichoptera | Polycentropodidae | *Plectrocnemia* |
| Polycentropodidae | Predator | Insecta | Trichoptera | Polycentropodidae | NA |
| *Tinodes* | Omnivore | Insecta | Trichoptera | Psychomyiidae | *Tinodes* |
| Psychomyiidae | Herbivore | Insecta | Trichoptera | Psychomyiidae | NA |
| *Rhyacophila* | Predator | Insecta | Trichoptera | Rhyacophilidae | *Rhyacophila* |
| *Sericostoma* | Omnivore | Insecta | Trichoptera | Sericostomatidae | *Sericostoma* |
| Sericostomatidae | Omnivore | Insecta | Trichoptera | Sericostomatidae | NA |
| *Corophium* | Detritivore | Malacostraca | Amphipoda | Corophiidae | *Corophium* |
| *Echinogammarus* | Omnivore | Malacostraca | Amphipoda | Gammaridae | *Echinogammarus* |
| *Gammarus* | Omnivore | Malacostraca | Amphipoda | Gammaridae | *Gammarus* |
| Gammaridae | Omnivore | Malacostraca | Amphipoda | Gammaridae | NA |
| *Dikerogammarus* | Omnivore | Malacostraca | Amphipoda | Gammaridae | *Dikerogammarus* |
| *Atyaephyra* | Detritivore | Malacostraca | Decapoda | Atyidae | *Atyaephyra* |
| *Orconectes* | Omnivore | Malacostraca | Decapoda | Cambaridae | *Orconectes* |
| Cambaridae | Omnivore | Malacostraca | Decapoda | Cambaridae | NA |
| Asellidae | Detritivore | Malacostraca | Isopoda | Asellidae | NA |
| Oligochaeta | Detritivore | Oligochaeta | NA | NA | NA |
| Dendrocoelidae | Predator | Turbellaria | Tricladida | Dendrocoelidae | NA |
| Dugesiidae | Predator | Turbellaria | Tricladida | Dugesiidae | NA |
| Planariidae | Predator | Turbellaria | Tricladida | Planariidae | NA |
